# Correcting for Superficial Bias in 7T Gradient Echo fMRI

**DOI:** 10.1101/2020.11.20.392258

**Authors:** Pei Huang, Marta M. Correia, Catarina Rua, Christopher T. Rodgers, Richard N. Henson, Johan D. Carlin

## Abstract

The arrival of submillimetre ultra high-field fMRI makes it possible to compare activation profiles across cortical layers. However, the Blood Oxygenation Level Dependent (BOLD) signal measured by Gradient-Echo fMRI is biased towards superficial layers of the cortex, which is a serious confound for laminar analysis. Several univariate and multivariate analysis methods have been proposed to correct this bias. We compare these methods using computational simulations and example human 7T fMRI data from Regions-of-Interest (ROIs) during a visual attention paradigm. The simulations show that two methods - the ratio of ROI means across conditions and a novel application of Deming regression - offer the most robust correction for superficial bias. Deming regression has the additional advantage that it does not require that the conditions differ in their mean activation over voxels within an ROI. When applied to the example dataset, these methods suggest that attentional modulation of activation is similar across cortical layers within the ventral visual stream, despite a naïve activation-based analysis producing stronger modulation in superficial layers. Our study demonstrates that accurate correction of superficial bias is crucial to avoid drawing erroneous conclusions from laminar analyses of Gradient-Echo fMRI data.

## 2 Introduction

Different layers in the neocortex support different types of neural computations. For instance, different layers of the visual cortex are preferentially involved in feedforward versus feedback connectivity (Rockland, 2017; Rockland and Pandya, 1979), suggesting that they encode distinct “bottom-up” and “top-down” processes. However, because the cortical ribbon is only 2-3mm thick in the sensory cortices (Economo, 1929), conventional fMRI acquisitions with 2-3mm isotropic voxels cannot resolve these layers (Dumoulin et al., 2017). Thus, most previous investigations of laminar organization involved invasive measurements that are generally unavailable in humans (Takahashi et al., 2016; Van Kerkoerle et al., 2017).

Recent advancement in MRI scanners, however, has changed the landscape. With higher field strengths (e.g, 7T), and in turn higher signal-to-noise ratios, human scanners are able to acquire data at submillimetre resolution, and thereby offer layer-specific or “laminar fMRI”. Recent 7T fMRI studies (e.g., Kok et al., 2016; Lawrence et al., 2019; Muckli et al., 2015; Polimeni et al., 2010) have suggested that top-down modulation of neural activity (for example, by attention or expectation) occurs in specific layers, though there is a disagreement about which layers. For example, Kok et al. (2016) used the Kanizsa triangle illusion to study attentional effects. By restricting their analysis to an ROI that responded to the part of the visual field containing illusory contours but no actual stimulus input, they observed evidence of strongest top-down feedback effects in the activation of deep layers of visual cortex. Lawrence et al. (2019) utilized visual gratings and manipulated both bottom-up effects of visual contrast and top-down effects of spatial attention. While they found bottom-up effects on activation were strongest in the middle layer, their top-down effects were strongest in the superficial layers, contrary to Kok et al. (2016). Muckli et al. (2015) used partially occluded images, where part of a visual scene was replaced with a blank square. They again focused on an ROI that responded to the occluded region of the visual field, but rather than measuring the mean activation of that ROI, they attempted to decode the pattern of activity over all voxels within that ROI. While decoding of bottom-up information (in unoccluded images) was above chance and relatively constant across layers, they found that above-chance decoding of top-down information (in partially occluded images) only occurred in superficial layers. These discrepancies in layer selectivity could reflect a dissociation between superficial and deep layers in terms of the type of feedback they receive (depending on the precise paradigm), or could arise due to different analysis methods, which may result in different sensitivity to effects in particular layers. This highlights the importance of examining the assumptions behind the acquisition and analysis methods that researchers use.

In terms of acquisition, most laminar fMRI studies, including the ones cited in the above paragraph, have used Gradient echo (GE) MRI sequences that are sensitive to the Blood Oxygenation-Level Dependent (BOLD) signal (Olman and Yacoub, 2011), but alternative sequences are also increasingly popular (Yacoub et al., 2013). The BOLD-GE sequences tend to provide good sensitivity to functional changes, but are also susceptible to signal artifacts, particularly from the large draining veins on the cortical surface (Boxerman et al., 1995; Rua et al., 2017). In addition, there is a confound of superficial bias, where larger signals are observed in the superficial layers relative to the deep layers. This superficial bias has two main contributory factors: a) the presence of draining veins, in which deoxygenated blood from the deep layers flows towards the superficial layers, resulting in an artificial increase in fMRI signal in the superficial layers that is not reflective of the underlying neural activation, and b) variations in baseline cerebral blood volume (CBV) and relaxation parameters across different cortical depths. The net effect is that BOLD-GE sequences had lower specificity and a bias towards higher signal in superficial layers, which complicates comparisons of task-related activation across layers.

An alternative class of sequences measure BOLD with spin echo (SE), or gradient and spin echo (GRASE) (Feinberg et al., 2015). These sequences are less prone to superficial bias but have lower functional contrast to noise ratios (Beckett et al., 2019). A final class of sequences substitutes BOLD for alternative metrics of neural activation. Cerebral blood volume (CBV) can be measured with vascular space occupancy (VASO) sequences (Huber et al., 2017b; Lu et al., 2013), and cerebral blood flow (CBF) can be measured using arterial spin labelling (ASL) sequences (Huber et al., 2017b; Kashyap et al., 2019; Petcharunpaisan, 2010). While these blood flow-based sequences are able to remove the spatial blurring due to draining veins, they come with drawbacks relative to GE-BOLD MRI, such as the largest CBV changes being localized in the arteries and potential dilation retrogradely in the upper layers relative to the location of neuronal activation (Uludağ and Blinder, 2018). Similar to GRASE, these methods also tend to have less sensitivity as a trade-off for their higher specificity (Huber et al., 2017b). Finally, VASO and ASL generally have longer TRs than GE and so are generally restricted to a smaller field of view (Huber et al., 2017a).

These limitations of alternative sequences have resulted in continued popularity for BOLD-GE laminar fMRI (Hollander et al., 2020; Kok et al., 2016; Lawrence et al., 2019; Liu et al., 2020). There is instead increasing interest in using post-processing analysis techniques to remove the bias in the BOLD signal towards superficial layers (Fracasso et al., 2018; Polimeni et al., 2010; Yacoub et al., 2013), thus mitigating the main drawback of BOLD-GE sequences.

There have been many different approaches to characterize and correct for this superficial bias, although few investigators have systematically explored whether different superficial bias-correction methods are expected to successfully correct bias under a particular model. In terms of bias models, (Huber, 2020) distinguishes three general classes of models: the linear-offset model, the multiplicative model and the leakage model. The multiplicative model seeks to emulate the effect of variations in baseline parameters across cortical depths, which has a multiplicative effect on the fMRI signal. Meanwhile, the leakage model attempts to capture the effect of draining veins on the fMRI data as it simulates the propagation of signal from the deep layers to the superficial layers, mimicking the effect of draining veins carrying oxygenated blood towards the superficial layers. In terms of methods that seek to correct bias in empirical studies, proposed solutions include Z-scoring timecourses (Lawrence et al., 2019), L2 normalization of beta estimates across conditions (Kay et al., 2019), taking the ratio of activations in two experimental conditions (Kashyap et al., 2017; Liu et al., 2020) and decoding using multi-voxel classification techniques (Hollander et al., 2020; Muckli et al., 2015).

Here, we explore how six of these bias-correction methods perform under a simple multiplicative model applied to a visual attention paradigm. Specifically, we simulated data with or without a (nonlinear) laminar profile, and compared how different bias-correction methods perform in recovering these ground truth profiles in the context of a superficial bias nuisance effect. We also introduce a new application of Deming regression to this problem, and found that this bias-correction method outperformed other commonly used approaches when conditions do not differ in regional-mean offset, while performing comparably to a region-mean ratio metric when conditions do differ. By introducing a method that provides robust correction for superficial bias, future studies will be able to benefit from the high sensitivity of BOLD-GE sequences for laminar fMRI.

## 3 Methods

Since the simulations are matched to the design of the example dataset, we describe this experiment first.

### 3.1 Experimental Design

We designed a block-based sustained attention paradigm. On each trial, participants performed a same-different judgment of two images that were presented at diagonally-offset locations around a central fixation cross. We varied the attended location (positive diagonal vs negative diagonal of a 2×2 stimulus array around the fixation cross) and stimulus category (face or house images) between blocks of 10 trials. Each run comprised 20 blocks in total: attending to houses along the positive diagonal (H^45^), attending to houses along the negative diagonal (H^135^), attending to faces along the positive diagonal (F^45^) and attending to faces along the negative diagonal (F^135^). The conditions were presented in a sequence that was randomised separately for each run.

Between runs, we also manipulated the presence of a second pair of distractor image sequences, which, when present, were located at the opposite location and drawn from the opposite stimulus category. For both simulation and our fMRI acquisition, there were a total of eight runs: four with distractors present (TaskD+) and four with distractors absent (TaskD-). For the fMRI acquisition, we alternated the presence or absence of distractors between runs. In the simulation, each run is detrended individually and hence, there is no overarching effect due to the order of runs.

We attribute any selectivity for category or location when the distractor is present (TaskD+) to attentional selection, because both locations and both categories are present on each block, which provides complete matching of the conditions at the level of bottom-up stimulus-driven responses. By contrast, location and category selectivity during the no distractor (Task D−) context could also reflect stimulus-driven effects of the stimulated visual quadrants or the presented object category. Some of the bias-correction methods we explore below are based on the notion that the no distractor (Task D−) context provides a control condition that can be used to correct any superficial bias in the estimates for attentional selectivity when the distractor is present (Task D+). We will expand on this idea below.

#### 3.1.1 Stimulus and Procedure

All stimuli were created using Matlab (2009a, The MathWorks, Natwick, MA, USA) and presented in the scanner using Presentation (v17.2). For the main experiment, the category stimuli were presented in a circular patch at four locations, diagonally from the fixation cross at 45°, 135°, 225° and 315° respectively and spanning 0.16°-2.42° eccentricity. The fixation cross was shifted up from the centre of the screen by 2° visual angle due to visual obstruction of the lower segment of the screen by the head coil. There were a total of 20 face images and 20 house images for each category. All images were presented in greyscale and histogram-matched to equate luminance and root mean squared contrast. Thus, any category selectivity cannot be attributed to differences in brightness or contrast between the categories.

At the start of each block, two white dots (0.10° eccentricity) appeared for 350ms indicating the pair of patches (either 45° and 225° [indicated by ^45^] or 135° and 315° [indicated by ^135^]) to which the participant should attend. This was followed by 550ms of fixation. The stimuli then appeared for 950ms, during which the participant was required to perform a same-different judgement on the two attended stimuli, followed by 550ms of fixation. The two stimuli were identical on 50% of trials. This stimulus-fixation trial was repeated 10 times within each block. Between blocks, there was a rest block of fixation with a duration of 1560ms (1260ms for the two participants with TR=2440ms, see Section 3.2 for more details). The duration was chosen to ensure that the start of each block was in sync with the start of a volume acquisition. In the distractor-absent condition, the display consisted of two stimuli from the attended category in the attended locations, while in the distractor-present condition, two additional stimuli from the other category appeared in the non-attended locations.

#### 3.1.2 Stimulus Design for localizer (fMRI)

The localizer session comprised of four runs of a category-selective localizer.

The category-selective localizer task comprised 15-second blocked presentations of sequences of faces, scenes, objects, scrambled objects and fixation, with 1s fixation between blocks. Each of these five block types appeared in a random order in each run. There were 20 blocks per run (four presentations of each block type). Within each block, 25 randomly selected stimuli from the current category were presented consecutively for 800ms each. Participants performed a 1-back matching task while fixating on a black dot in the middle of the screen.

We used the categorical localizer data to define the following ROIs according to conventional functional criteria (see below for details): occipital face area (OFA), fusiform face area (FFA), scene-selective transverse occipital sulcus (TOS), and parahippocampal place area (PPA).

### 3.2 Data Acquisition

Participants provided informed consent under a procedure approved by the institution’s local ethics committee (Cambridge Psychology Research Ethics Committee). A total of six healthy participants were scanned (2 females, age range 20-41, 2 participants were authors of this study).

Prior to the experiment, participants underwent a separate session of behavioural training with eye-tracking using an SMI high speed eye tracker. The participant attempted the same task and received feedback on their fixation levels after each run. We recorded calibrated eye position during each block, and measured fixation stability as the difference in standard deviation along the attended and neglected axes over the block. We repeated the task until this fixation stability metric met a criterion level of under 0.5 degrees visual angle mean difference for two consecutive runs. As eye-tracking was not available in the 7T scanner, the behavioural training was important to ensure that participants were able to perform the task while maintaining fixation.

Participants contributed data over a total of two MRI sessions; one session of retinotopic and category localizers at 3T and one main experimental session at 7T. The 3T data were acquired on a Siemens 3T Prisma-Fit scanner using a standard 32-channel head coil, while the 7T data were acquired on a Siemens 7T Terra scanner using the Nova Medical 1Tx/32Rx head coil. At the start of each session, we also acquired a MPRAGE (3T) or MP2RAGE (7T) (Marques et al., 2010) structural that was used for coregistration across sessions. Participants maintained fixation on a cross in the middle of the screen throughout, and fixation accuracy was verified using the 50Hz SMI MRI eye tracker system at 3T.

For 3T, the MPRAGE parameters were as follows: TR = 2,250 ms, TE = 3.02 ms, TI = 900 ms, GRAPPA = 2, FOV = 256 mm*256 mm*192 mm, Matrix size = 256*256*192, FA = 9°, ToA =~ 5 min and the EPI parameters were as follows: 3mm isotropic voxels, TR=2000ms, TE=30ms, FA=78°, Matrix size = 64*64*32, ToA=~11mins.

For the 7T data, the MP2RAGE parameters were as follows: TR = 4,300ms, TE = 1.99 ms, TI 1 = 840 ms, TI 2 = 2370ms, GRAPPA = 3, Matrix size = 320*320*224, FA 1 = 5 °, FA 2 = 6 ° and the EPI parameters were as follows: 0.8mm isotropic voxels, TR=2390ms (2440ms for two participants), TE= 24ms (24.4ms for two participants), FA=80°, GRAPPA = 3, Matrix size = 200*168*84, ToA=~11mins. The TR and TE were slightly longer for two participants due to the peripheral nerve stimulation threshold being exceeded in the 7T scanner.

### 3.3 Data Analysis

#### 3.3.1 MRI Data Pre-processing

For the functional volumes acquired on the 3T scanner, the images underwent slice time correction and rigid body realignment using SPM12 (www.fil.ion.ucl.ac.uk/spm). For the volumes acquired on the 7T scanner, the images first underwent slice time correction using SPM12, and then TOPUP (Andersson et al., 2003) in FSL (Niazy et al., 2004) was applied to estimate the susceptibility-induced distortions, using the first five volumes of each experimental run plus five additional volumes acquired before each run with the reverse phase-encoding direction. The resultant distortion correction was applied to the entire run. Each fMRI volume was then individually realigned to the structural using Boundary-Based Registration (BBR) (Greve and Fischl, 2009). In doing so, we use the structural as a reference to ensure that the volumes are coregistered with each other, which we previously showed to provide better realignment within and across runs for high-resolution fMRI data compared to standard methods that separate motion correction and co-registration (Huang et al., 2020).

#### 3.3.2 Regions of interest

For the categorical ROIs, activation t-maps where obtained using SPM12 by fitting a GLM to the fMRI data from the categorical localizer runs. The face-selective areas (FFA and OFA) were obtained from the t-score map from subtracting the object conditions from the face conditions. Similarly, the scene-selective areas (TOS and PPA) were obtained from the t-score map from subtracting the object conditions from the scene conditions. For each ROI, we identified the peak voxel in the expected anatomical location in a statistical map thresholded at p < 0.05 (uncorrected). We then grew the complete ROI by adding the contiguous voxel that was most selective for the localizing contrast iteratively until the ROI comprised 100 voxels (for implementation see https://github.com/jooh/roitools/blob/master/spm2roi.m).

To improve the stability of the estimates and our sensitivity to any variations in the data across layers, we combined the TOS, PPA, OFA and FFA regions to generate a pooled category-selective ROI. This approach is further justified in Section 4.2. Note that we flipped the sign of the contrast vector to align with the region’s expected preference (e.g. F^45^+F^135^-H^45^-H^135^ for FFA and H^45^+H^135^-F^45^-F^135^ for PPA). All subsequent analyses concern this pooled category-selective ROI unless specifically noted otherwise.

We coregistered the 3T localizer session to the main experiment 7T session. This was done by first coregistering the 3T functional images to the 3T structural image using the SPM coreg function. The 3T structural image was then coregistered to the 7T structural image, again using the SPM coreg function. The transformations from both coregistration steps were then applied to the ROI mask images. No further transformation was necessary since the BBR realignment process realigns the functional 7T images to the 7T structural image.

#### 3.3.3 Cortical-depth definition

The GM-WM (Grey Matter-White Matter) and GM-CSF (Grey Matter-Cerebrospinal fluid) boundaries were obtained from the Freesurfer’s reconstruction of each participant’s 7T structural image (Fischl et al., 2002). These boundaries were visually inspected to ensure that the segmentation was accurate. In cases of poor segmentation, the realignment between the structural and the Freesurfer segmentation template was manually adjusted prior to repeating the Freesurfer reconstruction. The boundaries were exported to CBStools (Bazin et al., 2012) and used to generate three equivolume (Leprince et al., 2015) segmentations of the GM. Each GM voxel was then assigned to one of the three layers using a winner-takes-all approach, in which the voxel is assigned to the layer with which it has the largest overlap.

### 3.4 Computational Simulations

Our computational simulations were carried out in Matlab (2019a, The MathWorks, Natwick, MA, USA); the code is available on github (https://github.com/MRC-CBU/LaminafMRIsimulations/tree/1.0).

For clarity, we describe our simulations in terms of categorical attentional selectivity for a face-selective area such as the fusiform face area (FFA). However, the model is a general account for how fMRI responses arise as a function of neural modulations of interest and layer bias of no interest, so is applicable to other scenarios.

We model the effects of attention on neuronal responses, and how such modulations manifest in fMRI activations. In our model, the signal component in *N*=*2500* fMRI voxels is expressed as a sum over neuronal populations that are purely selective for houses or for faces. Neurons fire at rate *f*=1 when their preferred category is present in the stimulus display, and *f*=0 to the presence of the non-preferred category. We generate the number of neurons *n* that prefer each stimulus category for each voxel by sampling a half-normal distribution, and control category selectivity by varying the standard deviation of the distribution over categories. For an ROI that prefers faces, we set the standard deviation of that distribution to 1.45 for faces and 0.7 for houses; for an un-selective ROI (see Section 0), we equate the standard deviations to 0.7 for faces and houses. Although selectivity at the voxel level in this formulation arises from differences in the relative frequency of face and house neurons, one could also interpret these standard deviation parameters as controlling the firing rates of equally-sized populations.

In our model, attention operates as a scale factor *a* that increases the responses of neurons coding the attended category, while leaving the unattended category unaffected. We simulate layer differences in the strength of attentional modulation by varying this *a* parameter. The signal component of the response *R* at voxel *v* to the category *c* can be expressed as

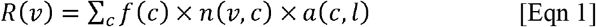

We modelled the neural response *R(v)* for the duration of each block, based on the timings of conditions in our experimental paradigm, and convolved these responses with a canonical hemodynamic response function to generate a timeseries for each voxel, *h(v,t)*.

We added two sources of noise: A physiological noise source, which has a low-dimensional structure over voxels and which scales with layer depth, and a white thermal noise source, which is constant in magnitude over the simulated volume. The rationale for the physiological noise is to model known low-dimensional fMRI noise processes such as heartbeat, respiration and residual head motion (Liu, 2016), which voxels exhibit to different extents. Formally, the physiological noise, *E*_*p*_, was modeled by generating 20 independent Gaussian noise vectors for the entire timecourse and projecting a randomly weighted combination of the vectors onto each voxel (Huang et al., 2018). This physiological noise was scaled to match a random zero-mean Gaussian distribution with standard deviation of σ_*p*_. The resulting timeseries was then multiplied by a scalar *L*_*Bias*_ to capture the bias towards superficial layers. Lastly, we added thermal noise, *E*_*t*_, from zero-mean Gaussian with standard deviation of σ_*t*_, which is independent of superficial bias. This was done for all 2500 voxels to generate a simulated timecourse, *y(v,t),* for voxel v in layer l. Thus:

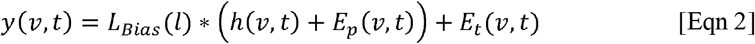

Having generated synthetic BOLD timeseries for each voxel, one can estimate the response to each condition by applying the General Linear Model (GLM), as is standard in fMRI analysis. To simulate the high-pass filtering commonly performed on real data to remove low-frequency noise in fMRI, first order sinusoidal and linear detrending was also performed. The GLM is based on the block timings and a canonical HRF, and produces a parameter estimate β for each condition and each voxel. The final contrast estimate (for a face-selective ROI) for the TaskD+ and TaskD-contrasts is then the subtraction of β for blocks when faces were unattended from that when blocks were attended.

To calculate the attentional contrast for our TaskD+ condition (when both positive and negative diagonals contain faces and houses, and the category being attended is equally often faces or houses), we take the simulated fMRI response during blocks where the participant is attending to the faces and subtract blocks where the participant is attending to the houses. The response for the TaskD+ contrast is therefore:

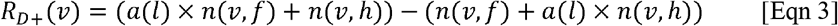

while in our TaskD- contrast, where faces or houses are only presented on the attended diagonal, the response is:

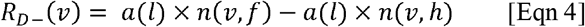

where *f* = face, *h* = house.

We explored a wide range of physiological and thermal noise parameters, and the final values of σ_*p*_ = 8 and σ_*t*_ = 15 were chosen by visually comparing scatterplots of TaskD+ against TaskD- for our simulated dataset against the experimental fMRI dataset for the pooled ROI (Figure 2). Furthermore, the combination of noise parameters and standard deviation values for the half-normal distributions were chosen such that the distribution of t-statistic for the simulated contrast was similar to the experimental dataset (see Supplementary Figure 1). Similarly, an attentional modulation factor of *a* = 3 was used to reflect the estimate generated from the experimental fMRI dataset. To simulate weaker attentional modulation in the middle layer, an attentional modulation factor of *a* = 2 was used for the middle layers.

**Figure 1:**
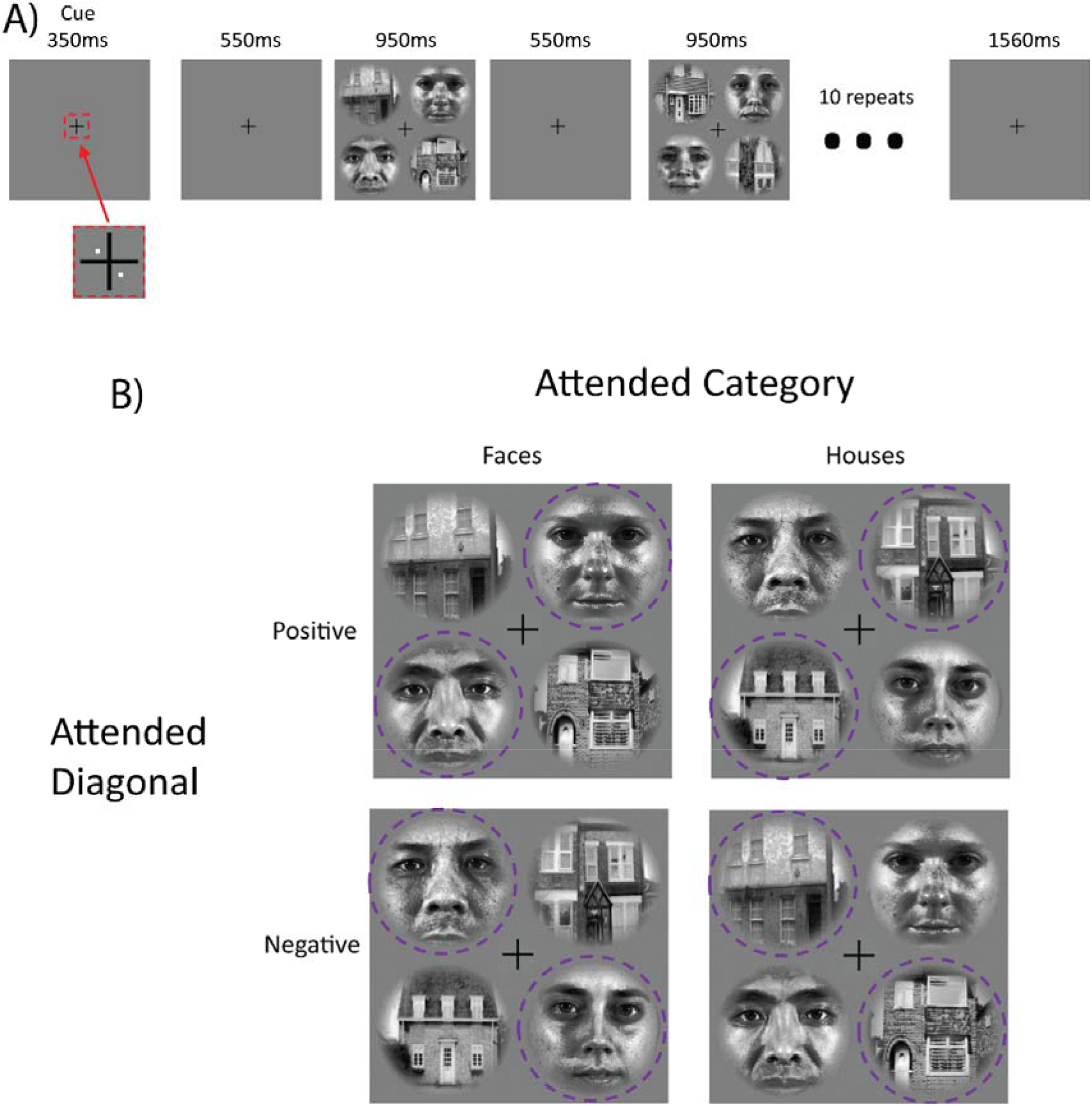
Panel A shows the experimental paradigm for each trial block. An initial pair of white dots cued participants to attend to a specific diagonal at the beginning of each task block. Ten image pairs from one category (faces or houses) appeared sequentially along the attended diagonal. In the “Task D+” distractor-present condition, image pairs from the other category appeared along the opposite diagonal (as shown here); in the “Task D- “ distractor-absent condition (not shown here), no stimuli appeared in the opposite diagonal. Participants decided whether the two stimuli on the attended diagonal were ‘same’ or ‘different’ (examples shown here are ‘different’); 50% of trials involved the same stimuli. Panel B illustrates the 4 main stimulus conditions. The purple dotted circle indicates the attended regions and was not present for the participant.

**Figure 2:**
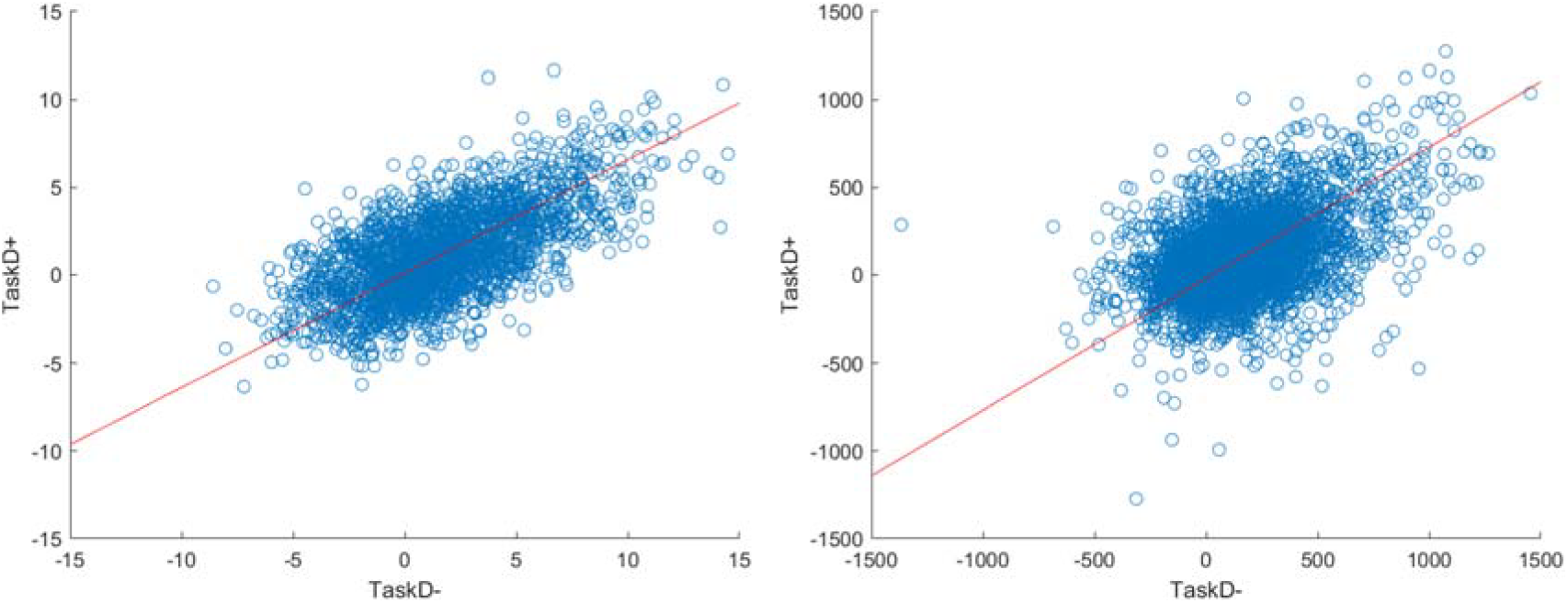
Scatterplots of the voxel responses (contrast estimates from the GLM) to TaskD+ against TaskD- comparing the simulated data with a = 3 (Panel A) against the real 7T data (Panel B). The scatterplots look similar, suggesting that the simulation is capturing the behaviour of real voxels. Note that the methods we evaluate are invariant to the absolute scale of the response, so these do not require matching; rather, we are only interested in the relation of the values across task types, i.e. the gradient of the best fit line, which is comparable across simulated and real data.

The six attentional modulation metrics were evaluated using simulated datasets with a V-shaped attentional modulation profile across layers of *a*(*l* = 1..3) = [3 2 3]. This profile indicates stronger attentional modulation in the superficial and deep layers, as might be expected from stronger feedback connectivity in these layers (Rockland, 2017; Rockland and Pandya, 1979). We repeated the simulation 10,000 times to calculate the accuracy and reliability of the six attentional modulation metrics.

### 3.5 Attentional Modulation Metrics

We were interested in validating the accuracy and reliability of the following six metrics of attentional modulation, which were applied to both the simulated and real data. The metrics were analysed in two ways. Firstly, the variation of each metric across layers was plotted and compared to the “ground truth” attentional modulation graph. Secondly, for each metric, we quantified the contributions of the superficial bias and attentional modulation towards the laminar profile. This was achieved by correlating the simulated data with either a [1 0 −1] contrast vector (for superficial bias) or [0.5 −1 0.5] (for attentional modulation) to determine the contributions of the two factors. The ideal laminar profile correlates with attentional modulation and not with superficial bias.

#### 3.5.1 Ratio metrics

The first group of attentional modulation metrics can be classified as ratio metrics; these metrics take the ratio between two experimental conditions (in this case, TaskD+ and TaskD-) to remove the superficial bias across layers. This method is motivated by the assumption that the baseline parameters, such as baseline blood volume and oxygen extraction fraction, have a multiplicative effect on BOLD sensitivity, as suggested in Kashyap et al. (2017). Thus, we can formulate the BOLD signal change as δ*S* = *L(l)*\**R*, where *L* is a function of the baseline physiological parameters that influence the BOLD response and *R* is the actual change in CBV and concentration of deoxygenated haemoglobin in response to neural activity. In our simulation, L_bias_(l) is used to model L(l). We assume that *L(l)* is constant within each cortical layer but varies across cortical layers, while *R* is the quantity of interest. Thus, by taking a ratio of the signal changes for two contrast estimates for each voxel, we can remove the dependence of the contrast on the baseline parameters:

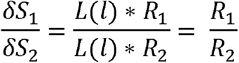

Note that while the attentional modulation, *a*, is defined to be greater than 1, the ratio metric of TaskD+/TaskD- in our simulation and experiment should always be less than 1. We refer to this ratio metric of TaskD+/TaskD- as selectivity, *S*, to differentiate it from the attentional modulation. The selectivity looks at the magnitude of top-down attentional modulation in TaskD+ as a fraction of both top-down attentional modulation and bottom-up stimulus activation in TaskD-. If we assume that our model is accurate, the relationship between selectivity and attentional modulation is:

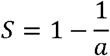

##### 3.5.1.1 Ratio of individual voxels (Voxel Ratio)

Given the above argument, one possible approach is to calculate the ratio of two values the GLM contrast for the TaskD+ condition relative to that for the TaskD- condition, for each voxel separately, and then average these ratios across voxels in a layer:

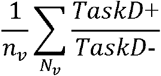

We refer to this metric as the “voxel ratio” throughout. However, this method has a high sensitivity to voxels where the TaskD-contrast is close to zero (e.g, because no attentional effect, or owing to random noise or fMRI susceptibility artifacts), which can produce extreme outliers. Assuming both TaskD+ and Task D-contrasts follow a Gaussian distribution, the resultant voxel ratio would have a heavy-tailed distribution, on which statistical tests are difficult. As such, we do not expect the voxel ratio to generate stable estimates but include it for completeness.

##### 3.5.1.2 Ratio of entire ROI (ROI Ratio)

To reduce the effect of outlier voxel values, one can first average (or sum) responses across voxels in a layer, before taking the ratio of the averages in each condition (cf. Guo et al., 2020):

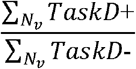

We refer to this metric as the “ROI ratio” for simplicity. A similar approach was adopted by Kashyap et al. (2017), where they compared the average of the activation peak against the average post-stimulus undershoot. However, by taking the average prior to the ratio, this method discards any pattern information that might be present within the voxels (we return to this point below).

##### 3.5.1.3 Deming Regression

Here, we chose not to average across voxels, but to estimate the gradient of the best-fitting line when regressing one condition against the other condition. Importantly, we used Deming regression to estimate this gradient, rather than ordinary least squares, because Deming regression accounts for errors in both variables (Adcock, 1878). A key benefit of Deming regression is that it does not require a regional-mean activation level difference between the two conditions, unlike the ROI ratio approach above. This makes the method potentially attractive for cases where voxels within an ROI have strong but opposing selectivity, as might for instance be expected in retinotopic maps of the visual field.

#### 3.5.2 Z-scoring of data

Rather than taking ratios or gradients of the estimated responses (i.e, contrasts of GLM Betas), another approach is to normalize the original data (*y(v,t)* in Eqn 2) before fitting the GLM, for example by using Z-scoring (Lawrence et al., 2019), in an attempt to equate the scaling across layers. Here, after Z-scoring the data and fitting the GLM, we simply calculated the contrast estimate for the TaskD+ condition. Z-scoring assumes that the noise and the signal of interest have a linear correlation. However, given that this is not the case in our simulation (physiological noise scales with the signal but thermal noise does not), we expect Z-scoring to perform worse than the ratio approaches.

#### 3.5.3 Multivoxel pattern analysis

The methods described above that first average over voxels within an ROI are only able to estimate a layer’s univariate activation. An alternative approach is to examine the multivariate information across voxels, e.g, by comparing the ability to classify whether faces or houses were attended. This was the approach taken by (Muckli et al., 2015). The logic underlying this approach is that cross-validation performance will be unaffected by superficial bias as long as the effect of interest and the noise scale together with overall signal magnitude. Like Z-scoring above, any additive (thermal) noise components will invalidate this assumption and reintroduce sensitivity to layer depth.

Here, we compared the ability of two methods of classification, Support Vector Machines classification (SVM) and Linear Discriminant Contrast (LDC) to discern between attending to houses vs faces in the TaskD+ condition. SVM is a robust method that has been widely used in fMRI analysis (Abdulkadir et al., 2013; Hoeft et al., 2011; Meier et al., 2012; Weygandt et al., 2012) and have also been used previously for laminar analysis (Muckli et al., 2015). LDC is often preferable for fMRI decoding since it is a continuous measure that does not exhibit ceiling effects (Huang et al., 2018; Kriegeskorte and Diedrichsen, 2016; Misaki et al., 2010; Yoon et al., 2012; Zhu et al., 2008).

Furthermore, unlike SVM, it does not require multiple instances for effective training, and therefore avoids issues to do with small numbers and temporal autocorrelation of single-trial/block estimates from the same run. Previous work demonstrated that LDC outperforms SVM in terms of detecting a difference in discriminability (Huang et al., 2018) and in terms of overall reproducibility of pairwise discriminants over splits in the data (Walther et al., 2016). Both SVM and LDC used leave-one-out cross-validation across the four runs of each task.

##### 3.5.3.1 SVM

We used the “fitcsvm” classifier in the Matlab Bioinformatics toolbox, with default settings (specifically, a linear kernel with C=1). To obtain multiple patterns for training, each block within a run was modelled with a separate regressor in a new GLM, producing 60 block estimates for training and 20 for testing on each fold of leave-one-run-out cross-validation. The final result is mean classification accuracy (% of test blocks correct).

##### 3.5.3.2 LDC

LDC is a continuous statistic related to Fisher’s linear discriminant. First, the training data is used to generate a set of weights to maximize the distance between the conditions of interest (e.g., houses vs faces). This set of weights is referred to as the discriminant. The LDC quantifies the difference between the two conditions in the testing data, measured on this discriminant.

As LDC does not require multiple training patterns, all blocks of the same type (within the training set) were modelled with a single regressor in the GLM. The contrast between attending to faces versus houses generates a distance metric, which is normalized using the sparse covariance matrix of the noise residuals (Ledoit and Wolf, 2003) to produce a weights vector. The dot product of the weights vector with the pairwise contrast estimate from the held-out test run produces the LDC test statistic. The final result is the average LDC across the four folds, normalized by the square root of the number of voxels (which differed by ROI in the real data).

## 4 Results

### 4.1 Computational Simulations

#### 4.1.1 Comparison of the six attentional modulation metrics

For our main simulations, we simulated an attentional modulation that is strongest in the superficial and deep layers and weaker in the middle layer. The simulated fMRI responses were then measured according to the six different metrics and compared against the ground truth profile to verify their accuracy and precision. Selectivity estimates using Deming regression, Voxel ratio and ROI ratio were plotted on the same axis as they generate commensurate estimates; the remaining three metrics were plotted on individual axes and scaled to best match the ground truth profile. In addition, we also plotted the raw contrast estimates and “ground truth” attentional modulation as a baseline comparison.

As can be seen in Figure 3, both Deming regression and ROI ratio were able to replicate the attentional modulation profile with high precision: The variability across iterations was low, as the percentile error bars illustrate. Though the voxel ratio recovered the V-shaped profile, it had very large error bars, owing to extremely high values for some voxels with a denominator (TaskD-) close to zero.

**Figure 3:**
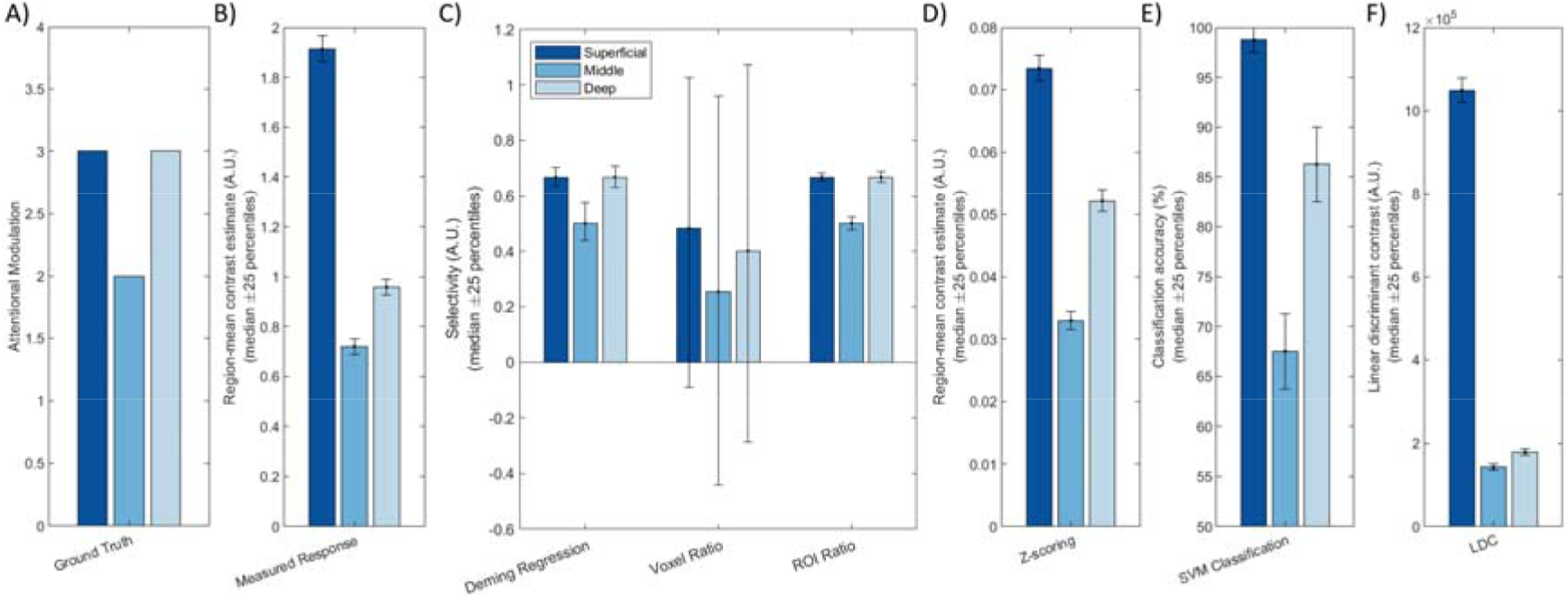
Simulation results showing A) the ground truth (neural response), B) the measured (BOLD) response, and C-F) the six attention metrics. Deming regression, Voxel Ratio and ROI Ratio were plotted on the same scale as they are commensurate; the other three metrics have different scales so were plotted individually. The different bar colours represent the different layers, with the dark blue representing the superficial layer and the light blue representing the deep layer. The bars indicate median performances over iterations of the simulation, while the error bars indicate the 25^th^ and 75^th^ percentile values.

Z-scoring the data before fitting the GLM produced a slanted V profile, indicating contributions from superficial bias as well as attentional modulation. The SVM and LDC classification methods produced similar slanted V profiles. The LDC method exhibited particularly dramatic superficial bias, with strong discriminant values for the superficial layer where the response magnitude was greatest.

#### 4.1.2 Quantifying the relative contributions of superficial bias and attentional modulation to laminar profile

We correlated the laminar profile of each metric (normalized by their mean) with either a [1 0 −1] or a [0.5 −1 0.5] vector to obtain a summary measure of the contributions of superficial bias and attentional modulation across layers. These summary measures indicate the reliability of any apparent layer differences and provide a means to compare the magnitude of attentional modulation and superficial bias effects within each metric.

As shown in Figure 4, both Deming regression and ROI ratio were highly correlated with the ground-truth attentional modulation effect, with almost zero contribution of superficial bias. Both metrics also exhibited low variability over iterations, as indicated by the 25-percentile error bars. By contrast, the voxel ratio was highly variable. While relatively stable over iterations, the Z-scoring metric was substantially influenced by the nuisance superficial bias effect. Similar effects were observed for the SVM metric. Finally, the LDC metric had the lowest variability over iterations, but exhibited the strongest influence of superficial bias out of all evaluated metrics.

**Figure 4:**
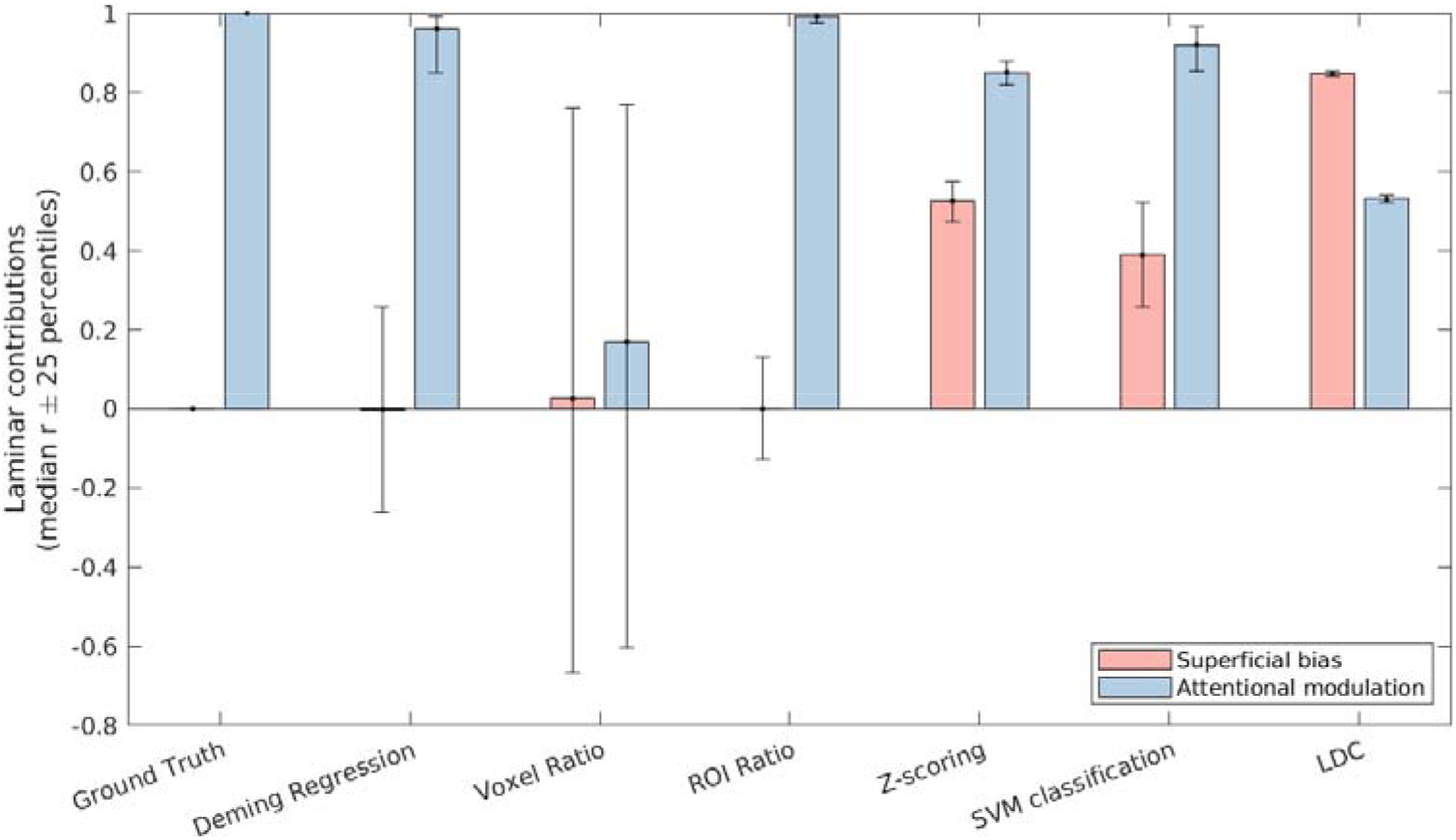
The contribution of superficial bias (pink bars) and attentional modulation (blue bars) to the laminar profile as recovered by the respective attentional metrics. The bars indicate the median correlation over iterations of the simulation, while the error bars indicate the 25^th^ and 75^th^ percentile values.

The SVM and LDC metrics exhibited distinct effects even though both are based on linear discriminants. We reasoned that this could reflect compressive effects of close-to-ceiling performance for the SVM (Figure 4E). This could obscure a superficial bias effect since the highest accuracy is in the superficial layer, where the bias is strongest. As a continuous metric, LDC does not exhibit ceiling effect. This can be demonstrated by repeating the simulation with increased noise levels (such as σ_*p*_ = 20 and σ_*t*_ = 30), under which the profiles for the SVM and LDC metrics were comparable, confirming that the apparent differences between the SVM and LDC metrics in Figures 4–5 reflect an SVM ceiling effect.

**Figure 5:**
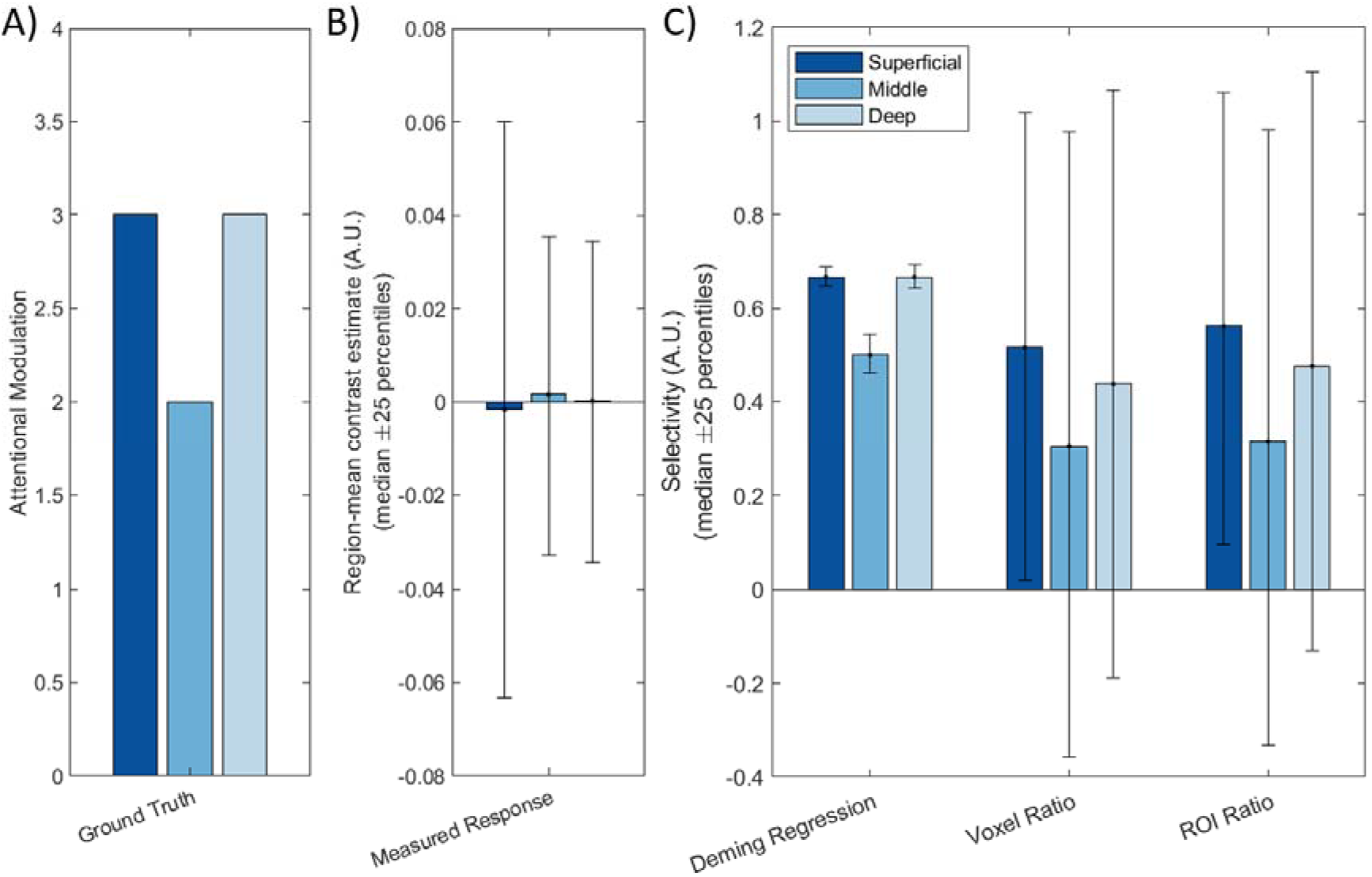
Comparing Deming regression and ratio of entire ROI with the ground truth in the absence of a global preference. The different bar colours represent the different layers, with the dark blue representing the superficial layer and the light blue representing the deep layer. The bars indicate the median effect over iterations of the simulation, while the error bars indicate the 25^th^ and 75^th^ percentile values.

#### 4.1.3 Simulating the effects of no region-mean preference

Both the Deming regression and ROI Ratio showed similar results in our initial simulations. However, Deming regression is potentially sensitive to local variations within an ROI, even if there is no global preference of that ROI. To illustrate this, we repeated the simulation with the same underlying [3 2 3] attentional modulation but removed the global preference for houses by sampling the density of house responsive cells and face responsive cells from the same distribution (folded normal distribution with mean 0 and standard deviation 0.7). Thus, each voxel has an equal chance of preferring faces or houses, with a resultant near-zero global preference for either house or faces (on average across simulations). As Figure 5B shows, there was no distinguishable variation in the measured response. However, Figure 6C shows that only Deming regression was able to recover the V-shape with sufficient sensitivity (i.e, relative to the error bars).

**Figure 6:**
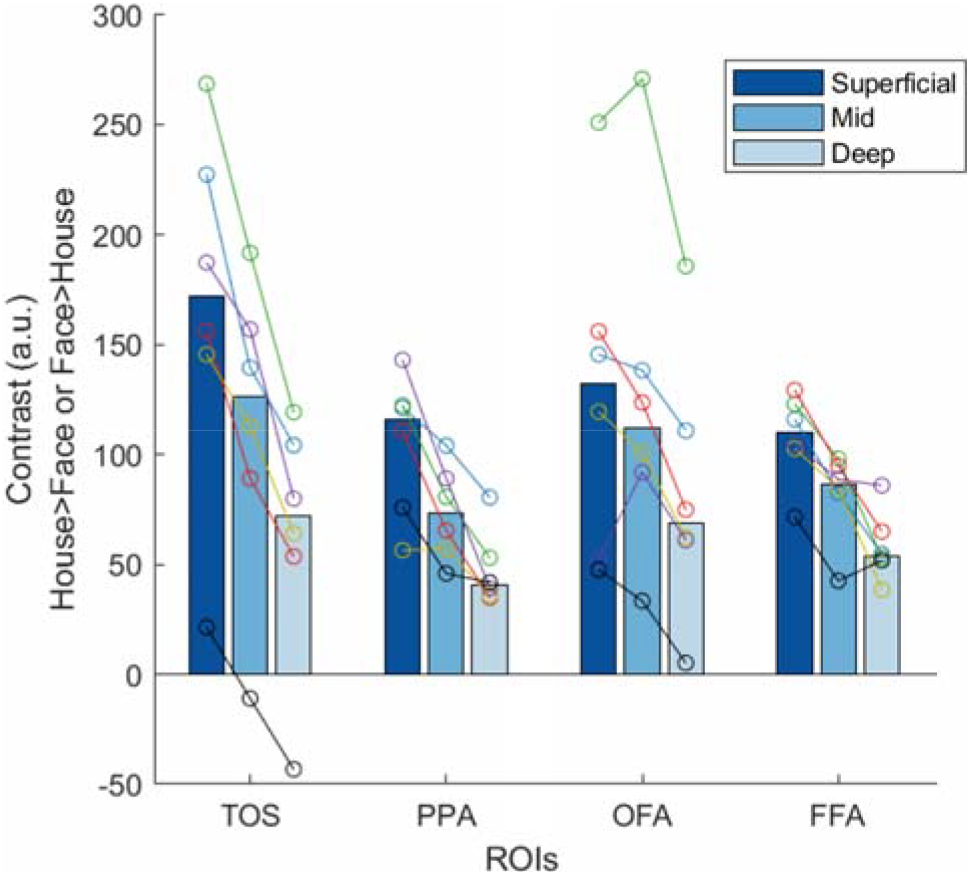
Plots of the categorical selectivity estimates obtained at 7T across different layers for the four different ROIs The bars represent the median of all six participants, while each colour in the overlaid circles represent an individual participant. The same colour represents the same participant throughout all plots.

It is interesting to note that, while the ROI ratio (like the Voxel ratio) is extremely noisy when there is no global preference, there is still a hint of a V-shape in the means in Figure 6C. We suspect this occurs because, while the overall preference averages to zero across all simulations, by chance, most individual simulations still have some global preference for either faces or houses. These small, but persistent, global preferences would still be sufficient for a ratio metric to recover the attentional modulation to some extent. However, as the percentile error bars illustrate, these ratio metrics are likely to be too variable to be useful when global preferences are near zero. This result demonstrates that Deming regression is the only method that can robustly detect layer specific modulations even in the absence of region-average differences.

### 4.2 Laminar analysis of real 7T data

Initial laminar analysis of the 7T data showed strong selectivity in the superficial layers with a constant decrease towards deeper layers, consistent across all ROIs (Figure 7). This is consistent with previous evidence for a superficial bias in GE sequences (Polimeni et al., 2010; Yacoub et al., 2013). We observed considerable variability across the six participants and four ROIs, so to generate a more robust estimate, we pooled voxels from TOS, PPA, OFA and FFA into a single pooled “category-selective” ROI. It can be seen that the pooled ROI exhibited substantially reduced between-participant variability (Figure 7A). In the following analyses, we focus on this pooled ROI, since we had no hypotheses concerning regional specificity of these effects.

**Figure 7:**
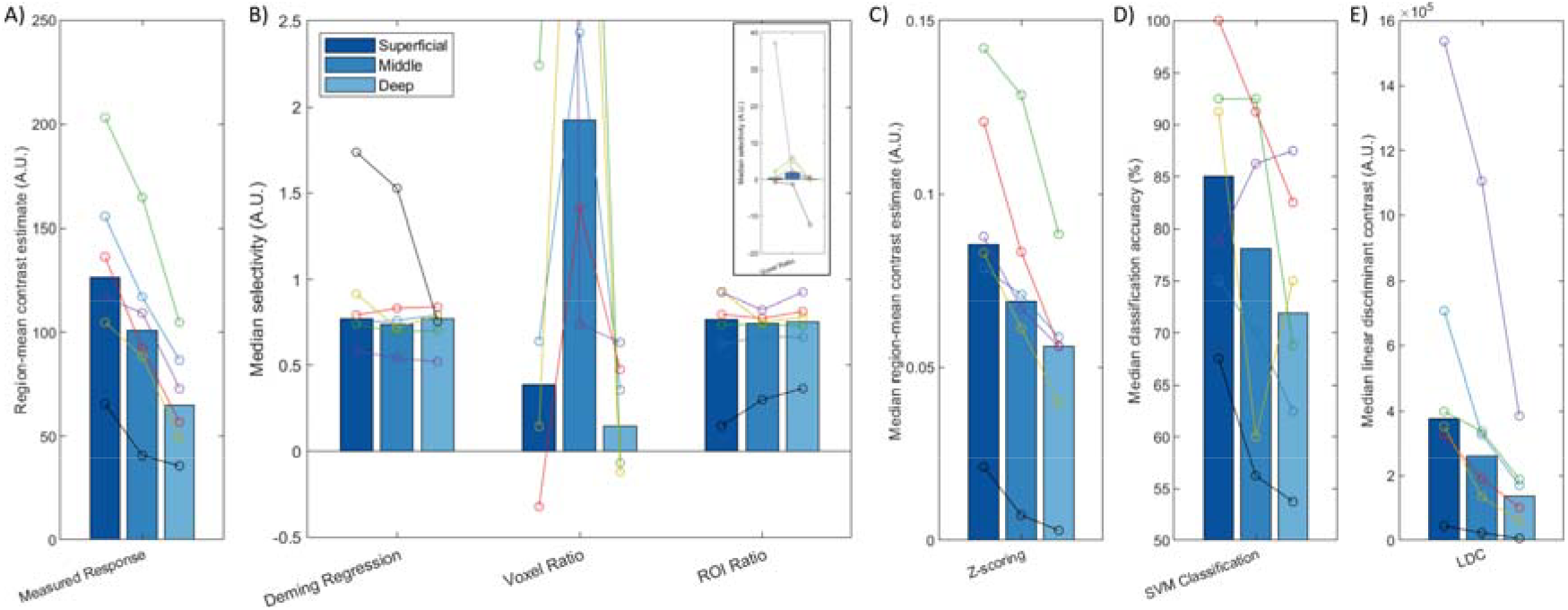
Plots of the measured response (Panel A) and attentional modulation metrics (Deming regression, Voxel ratio, ROI ratio, Z-scoring, SVM classification and LDC, Panels B-E) across different layers. These bars represent the median of all six participants, with each set of joint circles represent an individual subject. The same colour represents the same participant throughout all plots. The axis range of panel B has been restricted due to extreme outliers for the voxel ratio. The full range of this data is shown in the panel inset.

#### 4.2.1 Laminar analysis of 7T data using Attentional Modulation Metrics

The attentional modulation of the voxels as estimated with Deming regression was constant across layers (Figure 7B). Considered together with the simulation results, this indicates that the increased selectivity in superficial layers that we observed in Figure 6 can be explained by GE superficial bias. The ROI ratio metric demonstrated a similar lack of variation across layers, and comparable inter-subject variability. The voxel ratio was highly unstable across participants (see inset panel for full data range). The remaining three measures (Z-scoring, SVM classification and LDC) exhibited stronger effect in superficial layers (Figure 8C-E), consistent with the sensitivity to superficial bias that we observed in our simulations.

## 5 Discussion

A key challenge for interpreting layer analysis of high-resolution GE EPI fMRI data is that signal magnitude decreases with layer depth. This study investigates methods for correcting such superficial bias effects. We used computational simulations to evaluate the ability of six different metrics to recover an attentional modulation layer effect of interest in the presence of a superficial bias nuisance effect. Two of the evaluated methods were proposed by us (Deming regression, LDC) and the remaining have been previously used in the literature (Voxel and ROI Ratio, Z-scoring, SVM). Only the ROI Ratio and Deming Regression metrics were able to recover the attentional modulation layer effect accurately and precisely. While the remaining metrics were able to detect some underlying differences in attentional modulation across layers, they were either noisier than the aforementioned two metrics (in the case of the Voxel Ratio) or retained a substantial component of residual superficial bias (in the case of Z-scoring, or the multivoxel methods of SVM and LDC). Such partial correction for layer bias is particularly concerning because it might lead to mistaken inferences. Thus, the main contribution of our study is to demonstrate that Deming regression and ROI ratio are the most promising metrics for layer analysis of GE fMRI data.

Although the Deming regression and ROI ratio metrics performed similarly in the context of a region-mean activation-level difference between the conditions, we found that Deming Regression was better able to extract layer-profiles when we simulated a scenario where activation-level differences were near zero. This property may be useful in early visual areas such as V1, where an ROI might not show a global preference for two orientations of a visual stimulus, even though voxels within that ROI often show a local preference for one or other orientation (Alink et al., 2013; Kamitani and Tong, 2005). Conversely, our simulations suggest that the ROI ratio metric produces slightly less variable estimates when ROIs do exhibit strong activation-level differences. In summary, we recommend Deming regression as a general solution for correcting superficial bias in GE fMRI, although studies focused exclusively on effects carried by the regional mean may realise a small reduction in variability by adopting the ROI ratio metric instead.

The substantial residual superficial bias in the Z-scoring, SVM and LDC metrics can be explained by the characteristics of fMRI noise, which likely exhibit components that are both additive and multiplicative with respect to layer depth in GE fMRI data. For instance, Z-scoring assumes a linear relationship between the noise and the contrast of interest, and normalises the contrast by dividing it by an estimate of noise from the variance in the data. However, since our model includes both thermal (laminar invariant) and physiological (laminar dependent) noise sources, the variance in the data is not a perfect reflection of the superficial bias. Thus, Z-scoring is unable to fully correct the superficial bias in our simulations. Similarly, multivariate pattern analysis methods (SVM and LDC) fail to account for the superficial bias in the data because relatively greater contrast to noise ratios (due to superficial bias) leads to more robust contrast and better classification/LDC values. These metrics are only expected to correct layer bias successfully if all noise components scale with layer depth, a noise model that would be implausible for high-resolution fMRI where substantial thermal noise is inevitable.

Interestingly, LDC and SVM showed different sensitivities to superficial bias versus attentional modulation in the simulations (Figure 5). As we have observed previously (Huang et al., 2018), LDC results had smaller variability across iterations, reflecting greater stability in the estimates. However, LDC also demonstrated higher sensitivity to the laminar bias. We believe that this is primarily caused by the performance of the SVM classifier being close to 100% in the superficial layer. This ceiling effect results in a non-linear relationship between the functional activation and classification accuracy and partially masks the superficial bias. By contrast, LDC is expected to scale linearly with response magnitude (Arbuckle et al., 2019), which in this context means it is better able to capture the full extent of the superficial bias. A repeated analysis with higher noise levels confirmed that SVM and LDC metrics performed more similarly when the SVM classifier’s performance was brought down from ceiling levels.

Our numerical simulations (https://github.com/MRC-CBU/LaminafMRIsimulations) may be useful to test yet other approaches to correcting for superficial bias. Nonetheless, the code makes several simplifying assumptions for ease of calculation and generalization, which we list here for clarity. Firstly, we assumed that neurons are purely responsive to faces or houses only and that the voxel BOLD response is a simple sum of populations of face- and house-cells within the voxel. Secondly, we assumed that attention modulates responsive cells only by applying a gain factor and there is no constant additive component of attention. Thirdly, superficial bias was modelled as a gain factor on responses. Fourthly, we only modelled two sources of noise: at the level of the true hemodynamic response and at the level of measurement of that response, with only the former noise term being scaled by the laminar bias. While it is not obvious that changes in these assumptions would affect the current conclusions, this may be worth exploring in future simulations. Note also that we set the parameters of the simulation to match the ratio of activations in the two conditions from the voxels in real 7T data. While other parameter settings could produce different results, we think that it is unlikely for the other four metrics to outperform ROI ratio and Deming regression within the range of realistic parameters.

A final key limitation of our numerical simulations is that they are based on a multiplicative scaling model of superficial bias (Huber, 2020). While we believe this is more plausible than the linear-offset model discussed by the same reference, there are more sophisticated leakage models (e.g. Havlicek and Uludağ, 2020). Based on our anatomical understanding of the basis for superficial bias, it is likely that a more complex model consisting of both multiplicative effects (variations in BOLD response parameters across layers) and leakage effects (presence of draining veins across layers) is needed to fully capture the intricacies of the superficial bias. However, such a model would require more assumptions, and while they may be necessary to extract laminar profile for responses of a single condition versus baseline, we suspect that when the interest is in the difference between two or more conditions, taking an ROI ratio or estimating the slope of a Deming Regression is a simpler and effective way of removing the scaling component of the superficial bias effect. However it must be kept in mind that leakage effects are not corrected by these methods.

We also applied the six correction methods to the GE-fMRI 7T data acquired on the same paradigm. The uncorrected data showed the characteristic increase in BOLD signal towards superficial layers (Hollander et al., 2020; Kok et al., 2016). This superficial profile remained after Z-scoring or applying SVM or LDC, while the Voxel Ratio produced an extremely variable estimate with apparently-greatest modulation in the middle layer. However, the Deming Regression or ROI Ratio metrics produced a largely-flat profile across layers, with low variability across participants compared to the other metrics. These results are broadly consistent with the numerical simulations, and suggest that apparent modulations in selectivity across layers in the initial regional-mean contrasts can be explained in terms of superficial bias in the GE-fMRI signal, rather than a difference in attentional modulation as such. However, the small sample size of the initial study reported here prevents us from drawing strong inferences about the magnitude of layer-specific effects of attention in the population. We are planning a pre-registered replication of this work with a larger sample to more precisely estimate this experimental effect.

Our failure to observe a layer-specific effect of attention contrasts with other findings using similar manipulations of “top-down” processes (e.g., Muckli et al., 2015; Kok et al., 2016; Lawrence et al., 2019). These studies are quite heterogeneous in experimental design and have yet to be independently replicated, so completely consistent findings may not be expected. However, we note that some of these results were obtained using metrics that our numerical simulations suggest do not successfully correct superficial bias, which may provide a further explanation for any discrepancies. In particular, we believe that inadequate correction for superficial bias may explain the superficial profile reported by Lawrence et al. (2019) for spatial attention with a Z-scoring metric, and the superficial profile reported by Muckli et al. (2015) for SVM classification of visual content outside the voxelwise receptive field. Thus, it would be difficult to draw any conclusions as to the presence of any neuronal response differences across layers based on those results. By contrast, Kok et al. (2016) reported relatively stronger selectivity for illusory contours in deep layers, a result that runs opposite to the expected direction of a superficial bias effect. Such a pattern of result is not expected by superficial bias - instead the selectivity estimates in Kok et al would likely be stronger after correction for superficial bias. In general, we urge caution in interpreting GE fMRI layer analyses where superficial bias has not been appropriately corrected.

## 6 Conclusion

In this study, we demonstrate that Deming regression and ROI ratio can adjust for superficial bias across cortical layers. This is important because there is a growing wealth of high resolution, GE-fMRI data that can be used for laminar analysis, but a lack of consensus on the optimal way to analyse these data, which could partially explain the variance in conclusions drawn from these data. By proposing robust yet simple analysis methods to remove the superficial bias, such as the use of Deming Regression, we hope these will help reconcile results across different studies, and allow researchers to probe deeper into the function of neuronal activity in different cortical layers.

## 7 Acknowledgements

The authors are grateful to all the radiographers at MRC Cognition and Brain Sciences Unit, University of Cambridge and the Wolfson Brain Imaging Centre, University of Cambridge for their assistance in performing the scans and to the volunteers for participating in the experiment. The authors would also like to thank Kendrick N. Kay for helpful comments on a previous version of the manuscript. This research was funded by the National Institute for Health Research (NIHR) Biomedical Research Centre (BRC). The BRC is a partnership between Cambridge University Hospitals NHS Foundation Trust and the University of Cambridge, funded by the NIHR. The views expressed here are those of the authors are not necessarily those of the NIHR or the Department of Health and Social Care. CTR is funded by the Wellcome Trust and the Royal Society [098436/Z/12/B].

## References

Abdulkadir, A., Ronneberger, O., Christian Wolf, R., Pfleiderer, B., Saft, C., Kloppel, S., 2013. Functional and Structural MRI Biomarkers to Detect Pre-Clinical Neurodegeneration. Curr. Alzheimer Res. 10, 125–134. https://doi.org/10.2174/1567205011310020002

Abdulrahman, H., Henson, R.N., 2016. Effect of trial-to-trial variability on optimal event-related fMRI design: Implications for Beta-series correlation and multi-voxel pattern analysis. Neuroimage 125, 756–766. https://doi.org/10.1016/j.neuroimage.2015.11.009

Adcock, R.J., 1878. A Problem in Least Squares. Analyst 5, 53. https://doi.org/10.2307/2635758

Alink, A., Krugliak, A., Walther, A., Kriegeskorte, N., 2013. fMRI orientation decoding in V1 does not require global maps or globally coherent orientation stimuli. Front. Psychol. 4. https://doi.org/10.3389/fpsyg.2013.00493

Andersson, J.L.R., Skare, S., Ashburner, J., 2003. How to correct susceptibility distortions in spin-echo echo-planar images: Application to diffusion tensor imaging. Neuroimage 20, 870–888. https://doi.org/10.1016/S1053-8119(03)00336-7

Arbuckle, S.A., Yokoi, A., Pruszynski, J.A., Diedrichsen, J., 2019. Stability of representational geometry across a wide range of fMRI activity levels. Neuroimage 186, 155–163. https://doi.org/10.1016/j.neuroimage.2018.11.002

Bazin, P., Weiss, M., Dinse, J., Schaefer, A., Trampel, R., Turner, R., 2012. A computational pipeline for subject-specific, ultra-high resolution cortical analysis at 7 Tesla. 18th Annu. Meet. Organ. Hum. Brain Mapp. 883.

Beckett, A.J., Dadakova, T., Townsend, J., Huber, L., Park, S., Feinberg, D.A., 2019. Comparison of BOLD and CBV using 3D EPI and 3D GRASE for cortical layer fMRI at 7T. bioRxiv 778142. https://doi.org/10.1101/778142

Boxerman, J.L., Bandettini, P.A., Kwong, K.K., Baker, J.R., Davis, T.L., Rosen, B.R., Weisskoff, R.M., 1995. The intravascular contribution to fmri signal change: monte carlo modeling and diffusionlweighted studies in vivo. Magn. Reson. Med. 34, 4–10. https://doi.org/10.1002/mrm.1910340103

Dumoulin, S.O., Fracasso, A., van der Zwaag, W., Siero, J.C.W., Petridou, N., 2017. Ultra-high field MRI: Advancing systems neuroscience towards mesoscopic human brain function. Neuroimage 168, 345–357. https://doi.org/10.1016/j.neuroimage.2017.01.028

Economo, C. von, 1929. The Cytoarchitectonics of the Human Cerebral Cortex.

Feinberg, D.A., Vu, A.T., Goebel, R., Kemper, V.G., Poser, B.A., Yacoub, E., De Martino, F., 2015. Sub-millimeter T2 weighted fMRI at 7 T: comparison of 3D-GRASE and 2D SE-EPI. Front. Neurosci. 9. https://doi.org/10.3389/fnins.2015.00163

Fischl, B., Salat, D.H., Busa, E., Albert, M., Dieterich, M., Haselgrove, C., Van Der Kouwe, A., Killiany, R., Kennedy, D., Klaveness, S., Montillo, A., Makris, N., Rosen, B., Dale, A.M., 2002. Whole brain segmentation: Automated labeling of neuroanatomical structures in the human brain. Neuron 33, 341–355. https://doi.org/10.1016/S0896-6273(02)00569-X

Fracasso, A., Luijten, P.R., Dumoulin, S.O., Petridou, N., 2018. Laminar imaging of positive and negative BOLD in human visual cortex at 7 T. Neuroimage 164, 100–111. https://doi.org/10.1016/j.neuroimage.2017.02.038

Greve, D.N., Fischl, B., 2009. Accurate and robust brain image alignment using boundary-based registration. Neuroimage 48, 63–72. https://doi.org/10.1016/j.neuroimage.2009.06.060

Havlicek, M., Uludağ, K., 2020. A dynamical model of the laminar BOLD response. Neuroimage 204. https://doi.org/10.1016/j.neuroimage.2019.116209

Hoeft, F., McCandliss, B.D., Black, J.M., Gantman, A., Zakerani, N., Hulme, C., Lyytinen, H., Whitfield-Gabrieli, S., Glover, G.H., Reiss, A.L., Gabrieli, J.D.E., 2011. Neural systems predicting long-term outcome in dyslexia. Proc. Natl. Acad. Sci. 108, 361–366. https://doi.org/10.1073/pnas.1008950108

Hollander, G. De, Zwaag, W. Van Der, Qian, C., Zhang, P., Knapen, T., 2020. Ultra-high resolution fMRI reveals origins of feedforward and feedback activity within laminae of human ocular dominance columns. bioRxiv 1–41. https://doi.org/10.1101/2020.05.19.102186

Huang, P., Carlin, J.D., Alink, A., Kriegeskorte, N., Henson, R.N., Correia, M.M., 2018. Prospective motion correction improves the sensitivity of fMRI pattern decoding. Hum. Brain Mapp. 39, 4018–4031. https://doi.org/10.1002/hbm.24228

Huang, P., Carlin, J.D., Henson, R.N., Correia, M.M., 2020. Improved motion correction of submillimetre 7T fMRI time series with Boundary-Based Registration (BBR). Neuroimage 210. https://doi.org/10.1016/j.neuroimage.2020.116542

Huang, P., Kriegeskorte, N., Henson, R., Alink, A., Correia, M., 2017. Quantifying the effectiveness of prospective motion correction using a visual fMRI task, in: Proceedings of the 25th Scientific Meeting of ISMRM. p. 1276.

Huber, L., 2020. Removing unwanted venous signal from GE-BOLD maps: Overview of vein removal models and implementations in LAYNII [WWW Document]. URL https://layerfmri.com/2020/04/02/devein/

Huber, L., Ivanov, D., Hall, A., Guidi, M., Bandettini, P.A., Gonzalez-Castillo, J., Goense, J., Chen, G., Marrett, S., Poser, B.A., Stüber, C., Handwerker, D.A., Jangraw, D.C., 2017a. High-Resolution CBV-fMRI Allows Mapping of Laminar Activity and Connectivity of Cortical Input and Output in Human M1. Neuron 96, 1253-1263.e7. https://doi.org/10.1016/j.neuron.2017.11.005

Huber, L., Uludağ, K., Möller, H.E., 2017b. Non-BOLD contrast for laminar fMRI in humans: CBF, CBV, and CMRO2. Neuroimage. https://doi.org/10.1016/j.neuroimage.2017.07.041

Kamitani, Y., Tong, F., 2005. Decoding the visual and subjective contents of the human brain. Nat. Neurosci. 8, 679–685. https://doi.org/10.1038/nn1444

Kashyap, S., Ivanov, D., Havlicek, M., Poser, B., Uludag, K., 2019. Laminar CBF and BOLD fMRI in the human visual cortex using arterial spin labelling at 7T, in: Proceedings of the 27th Scientific Meeting of ISMRM. p. 609.

Kashyap, S., Ivanov, D., Havlicek, M., Poser, B.A., Uludağ, K., 2017. Impact of acquisition and analysis strategies on cortical depth-dependent fMRI. Neuroimage. https://doi.org/10.1016/j.neuroimage.2017.05.022

Kay, K., Jamison, K.W., Vizioli, L., Zhang, R., Margalit, E., Ugurbil, K., 2019. A critical assessment of data quality and venous effects in sub-millimeter fMRI. Neuroimage 189, 847–869. https://doi.org/10.1016/j.neuroimage.2019.02.006

Kok, P., Bains, L.J., Van Mourik, T., Norris, D.G., De Lange, F.P., 2016. Selective activation of the deep layers of the human primary visual cortex by top-down feedback. Curr. Biol. 26, 371–376. https://doi.org/10.1016/j.cub.2015.12.038

Kriegeskorte, N., Diedrichsen, J., 2016. Inferring brain-computational mechanisms with models of activity measurements. Philos. Trans. R. Soc. Lond. B. Biol. Sci. 371, 489–495. https://doi.org/10.1098/rstb.2016.0278

Lawrence, S.J., Norris, D.G., de Lange, F.P., 2019. Dissociable laminar profiles of concurrent bottom-up and top-down modulation in the human visual cortex. Elife 8. https://doi.org/10.7554/eLife.44422

Ledoit, O., Wolf, M., 2003. Improved estimation of the covariance matrix of stock returns with an application to portfolio selection. J. Empir. Financ. 10, 603–621. https://doi.org/10.1016/S0927-5398(03)00007-0

Leprince, Y., Poupon, F., Delzescaux, T., Hasboun, D., Poupon, C., Riviere, D., 2015. Combined Laplacian-equivolumic model for studying cortical lamination with ultra high field MRI (7 T). Proc. - Int. Symp. Biomed. Imaging 2015-July, 580–583. https://doi.org/10.1109/ISBI.2015.7163940

Liu, C., Guo, F., Qian, C., Zhang, Z., Sun, K., Wang, D.J., He, S., Zhang, P., 2020. Layer-dependent multiplicative effects of spatial attention on contrast responses in human early visual cortex. Prog. Neurobiol. 101897. https://doi.org/10.1016/j.pneurobio.2020.101897

Liu, T.T., 2016. Noise contributions to the fMRI signal: An overview. Neuroimage 143, 141–151. https://doi.org/10.1016/j.neuroimage.2016.09.008

Lu, H., Hua, J., van Zijl, P.C.M., 2013. Noninvasive functional imaging of cerebral blood volume with vascular-space-occupancy (VASO) MRI. NMR Biomed. 26, 932–948. https://doi.org/10.1002/nbm.2905

Marques, J.P., Kober, T., Krueger, G., van der Zwaag, W., Van de Moortele, P.F., Gruetter, R., 2010. MP2RAGE, a self bias-field corrected sequence for improved segmentation and T1-mapping at high field. Neuroimage 49, 1271–1281. https://doi.org/10.1016/j.neuroimage.2009.10.002

Meier, T.B., Desphande, A.S., Vergun, S., Nair, V.A., Song, J., Biswal, B.B., Meyerand, M.E., Birn, R.M., Prabhakaran, V., 2012. Support vector machine classification and characterization of age-related reorganization of functional brain networks. Neuroimage 60, 601–613. https://doi.org/10.1016/j.neuroimage.2011.12.052

Misaki, M., Kim, Y., Bandettini, P.A., Kriegeskorte, N., 2010. Comparison of multivariate classifiers and response normalizations for pattern-information fMRI. Neuroimage 53, 103–118. https://doi.org/10.1016/j.neuroimage.2010.05.051

Muckli, L., De Martino, F., Vizioli, L., Petro, L.S., Smith, F.W., Ugurbil, K., Goebel, R., Yacoub, E., 2015. Contextual Feedback to Superficial Layers of V1. Curr. Biol. 25, 2690–2695. https://doi.org/10.1016/j.cub.2015.08.057

Niazy, R.K., De Stefano, N., Beckmann, C.F., De Luca, M., Vickers, J., Matthews, P.M., Behrens, T.E.J., Saunders, J., Flitney, D.E., Jenkinson, M., Zhang, Y., Bannister, P.R., Woolrich, M.W., Smith, S.M., Brady, J.M., Johansen-Berg, H., Drobnjak, I., 2004. Advances in functional and structural MR image analysis and implementation as FSL. Neuroimage 23, S208–S219. https://doi.org/10.1016/j.neuroimage.2004.07.051

Olman, C.A., Yacoub, E., 2011. High-Field fMRI for Human Applications: An Overview of Spatial Resolution and Signal Specificity. Open Neuroimag. J. 5, 74–89. https://doi.org/10.2174/1874440001105010074

Petcharunpaisan, S., 2010. Arterial spin labeling in neuroimaging. World J. Radiol. 2, 384. https://doi.org/10.4329/wjr.v2.i10.384

Polimeni, J.R., Fischl, B., Greve, D.N., Wald, L.L., 2010. Laminar analysis of 7T BOLD using an imposed spatial activation pattern in human V1. Neuroimage 52, 1334–1346. https://doi.org/10.1016/j.neuroimage.2010.05.005

Rockland, K.S., 2017. What do we know about laminar connectivity? Neuroimage. https://doi.org/10.1016/j.neuroimage.2017.07.032

Rockland, K.S., Pandya, D.N., 1979. Laminar origins and terminations of cortical connections of the occipital lobe in the rhesus monkey. Brain Res. 179, 3–20. https://doi.org/10.1016/0006-8993(79)90485-2

Rua, C., Costagli, M., Symms, M.R., Biagi, L., Donatelli, G., Cosottini, M., Del Guerra, A., Tosetti, M., 2017. Characterization of high-resolution Gradient Echo and Spin Echo EPI for fMRI in the human visual cortex at 7 T. Magn. Reson. Imaging 40, 98–108. https://doi.org/10.1016/j.mri.2017.04.008

Takahashi, N., Oertner, T.G., Hegemann, P., Larkum, M.E., 2016. Active cortical dendrites modulate perception. Science (80-.). 354, 1587–1590. https://doi.org/10.1126/science.aah6066

Uludağ, K., Blinder, P., 2018. Linking brain vascular physiology to hemodynamic response in ultra-high field MRI. Neuroimage 168, 279–295. https://doi.org/10.1016/j.neuroimage.2017.02.063

Van Kerkoerle, T., Self, M.W., Roelfsema, P.R., 2017. Layer-specificity in the effects of attention and working memory on activity in primary visual cortex. Nat. Commun. 8. https://doi.org/10.1038/ncomms13804

Walther, A., Nili, H., Ejaz, N., Alink, A., Kriegeskorte, N., Diedrichsen, J., 2016. Reliability of dissimilarity measures for multi-voxel pattern analysis. Neuroimage 137, 188–200. https://doi.org/10.1016/j.neuroimage.2015.12.012

Weygandt, M., Blecker, C.R., Schäfer, A., Hackmack, K., Haynes, J.D., Vaitl, D., Stark, R., Schienle, A., 2012. FMRI pattern recognition in obsessive-compulsive disorder. Neuroimage 60, 1186–1193. https://doi.org/10.1016/j.neuroimage.2012.01.064

Yacoub, E., De Martino, F., Zimmermann, J., Ugurbil, K., Goebel, R., Muckli, L., Ugurbil, K., Yacoub, E., Goebel, R., 2013. Cortical Depth Dependent Functional Responses in Humans at 7T: Improved Specificity with 3D GRASE. PLoS One 8, e60514. https://doi.org/10.1371/journal.pone.0060514

Yoon, J.H., Nguyen, D. V., McVay, L.M., Deramo, P., Minzenberg, M.J., Ragland, J.D., Niendham, T., Solomon, M., Carter, C.S., 2012. Automated classification of fMRI during cognitive control identifies more severely disorganized subjects with schizophrenia. Schizophr. Res. 135, 28–33. https://doi.org/10.1016/j.schres.2012.01.001

Zhu, C.Z., Zang, Y.F., Cao, Q.J., Yan, C.G., He, Y., Jiang, T.Z., Sui, M.Q., Wang, Y.F., 2008. Fisher discriminative analysis of resting-state brain function for attention-deficit/hyperactivity disorder. Neuroimage 40, 110–120. https://doi.org/10.1016/j.neuroimage.2007.11.029

