## Supplementary Figure 1 for "Correcting for Superficial Bias in 7T Gradient Echo fMRI"

### Supplementary Data

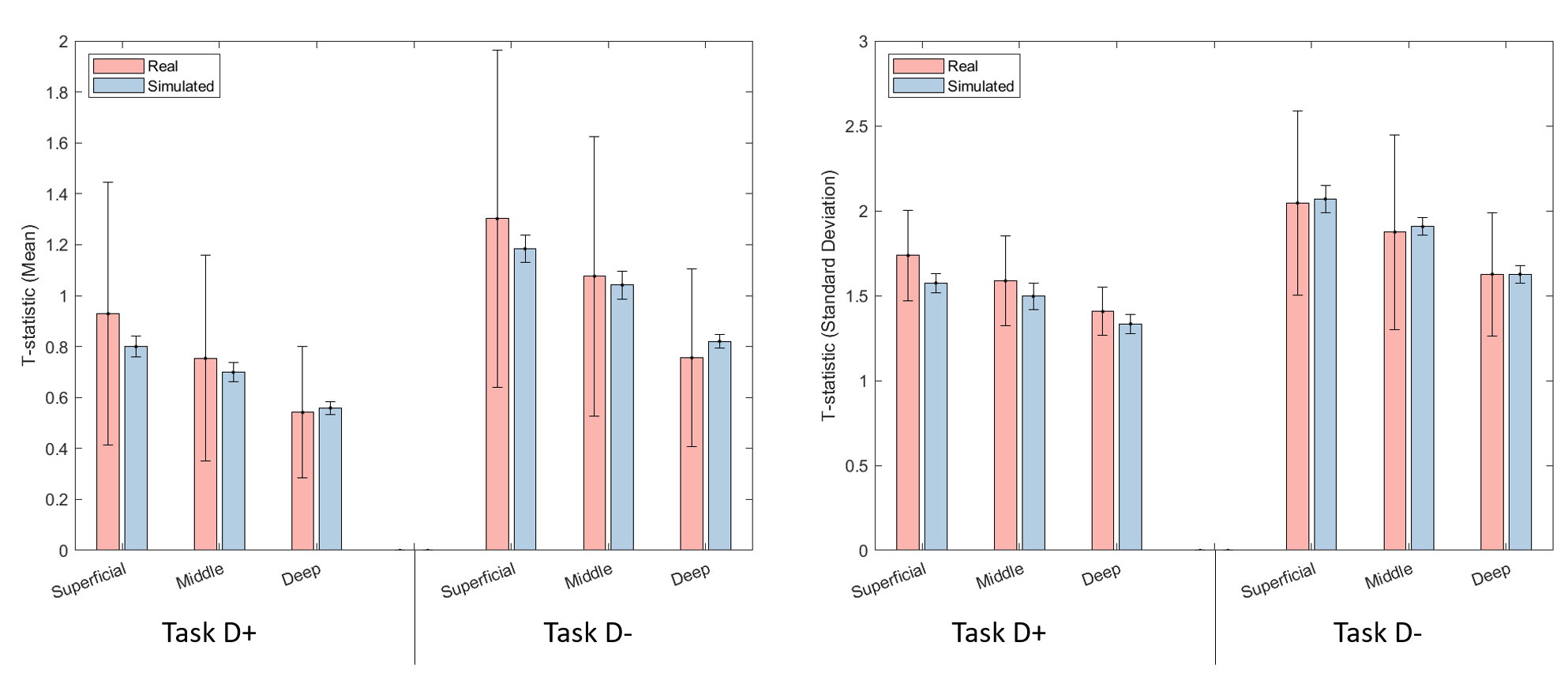

Supplementary Figure 1: Comparison of mean and standard deviation of t-statistic of the voxels in the ROI for real and simulated data. The error bars indicate the standard deviation of the summary metrics of the voxels across participants/iterations.
